# Meiosis and Kinetochore genes are used by cancer cells as genome destabilizers and transformation catalysts

**DOI:** 10.1101/826081

**Authors:** Roshina Thapa, Swetha Vasudevan, Mimi Abo-Ayoub Ashqar, Eli Reich, Nataly Kravchenko-Balasha, Michael Klutstein

## Abstract

Cancer cells have an altered transcriptome which contributes to their altered behaviors compared to normal cells. Indeed, many tumors express high levels of genes participating in meiosis or kinetochore biology, but the role of this high expression has not been fully elucidated. In this study we explore the relationship between this overexpression and genome instability and transformation capabilities of cancer cells. For this, we obtained expression data from 5 different cancer types which were analyzed using computational information-theoretic analysis. We were able to show that highly expressed meiotic/kinetochore genes were enriched in the altered gene expression subnetworks characterizing unstable cancer types with high chromosome instability (CIN). However, altered subnetworks found in the cancers with low CIN did not include meiotic and kinetochore genes. Representative gene candidates, found by the analysis to be correlated with a CIN phenotype, were further explored by transfecting genomically-stable (HCT116) and unstable (MCF7) cancer cell lines with vectors overexpressing those genes. This overexpression resulted in an increase in the numbers of abnormal cell divisions and defective spindle formations and in increased transformation properties in stable cancer HCT116 cells. Interestingly, the same properties were less affected by the overexpressed genes in the unstable MCF7 cancer cells. Our results indicate that overexpression of both meiosis and kinetochore genes is capable of driving genomic instability and cancer progression.

## Introduction

Cancer is a complex disease, characterized by numerous genomic aberrations and by dysregulation of gene expression. Along with overexpression of oncogenes and repression of tumor suppressors, tumors often express various tissue specific genes, not necessarily related to their primary tissue of origin [1–3]. In particular, cancer cells have been found to frequently express genes that are normally restricted to the testis. These genes can be referred to as cancer/testis antigens (CTA) [4]. Many CTA have been found to be involved in meiotic divisions, which occur in the testis and include processes with inherent genome instability. This property of the meiotic genes has led to the intriguing idea that the expression of CTA in tumors may drive genome and chromosome instability in those tumors [5,6].

Chromosome instability (CIN) is among the most important cancer hallmarks. CIN tumors, which have the propensity to constantly change their genome, have worse prognosis than non-CIN tumors [7,8]. Previous work shows that CIN tumors use several molecular mechanisms to achieve their instability, such as replication stress and modulation of the spindle assembly checkpoint [9–11]. Due to inherent functions of CTA genes involved in meiosis, that include mono-orientation of sister kinetochores and DNA double-strand break formation and repair, CTA have become prime candidates for initiating an additional mechanism involved in CIN [12–15].

Several small scale studies have already shown that a cohort of meiotic genes is expressed in different tumors [4], [6,16–19]. Importantly, a previous study has shown that overexpression of the meiotic cohesin Rec8 in mitotic fission yeast cells causes uniparental disomy of chromosomes and CIN in this organism [20].

Another emerging player in the generation of CIN in cancer is the kinetochore [21]. Kinetochores are protein complexes built on centromeres, the specialized loci on eukaryotic chromosomes, which play a key role in mediating chromosome segregation [21]. This is mainly achieved through the physical connection between microtubules and the centromeric DNA [22]. The balance between all the different kinetochore components is crucial for maintaining genome stability and correct ploidy. Under- or overexpression of different kinetochore components may lead to the formation of chromosomes with very little microtubule attachment, or on the contrary, too many microtubules binding to a chromosome [22–25]. Eventually this may lead to non-disjunction and aneuploidy [23–25]. Overexpression of specific kinetochore components such as the inner-centromere protein CENP-A (centromere specific ortholog of histone H3 serving as the structural basis of the kinetochore) leads to deposition of kinetochore components on additional loci in a chromosome already containing a centromere, and the formation of di-centric chromosomes, resulting in a breakage-fusion-bridge cycle of chromosomes and CIN [26]. On the other hand, insufficient CENP-A can result in senescence of cells and apoptosis [27,28]. Misregulation of kinetochore components has been observed in many tumors [29]. Alterations in the expression levels of kinetochore genes may also cause CIN in tumors, as well as affect the prognosis of specific patients and their response to therapy [29]. Despite all these studies the role that the kinetochores play as drivers of CIN during tumorigenesis is not fully understood.

To explore further the relationship between meiosis and kinetochore genes and genome instability we performed a large scale computational analysis of normal and cancer tissues which were obtained from breast, bladder, stomach, colorectal and cervical cancer and normal tissues, all from TCGA data (https://portal.gdc.cancer.gov/). We have demonstrated that tumors with high CIN harbored cancer-specific gene-gene correlation subnetworks with induced meiosis and kinetochore genes. Although tumors were heterogeneous and could be characterized by different altered gene expression subnetworks, meiosis and kinetochore altered transcripts could be found in various compositions in high CIN tumors but not in low CIN patients within the same type of cancer (see **Fig. 1**).

**Figure 1.**
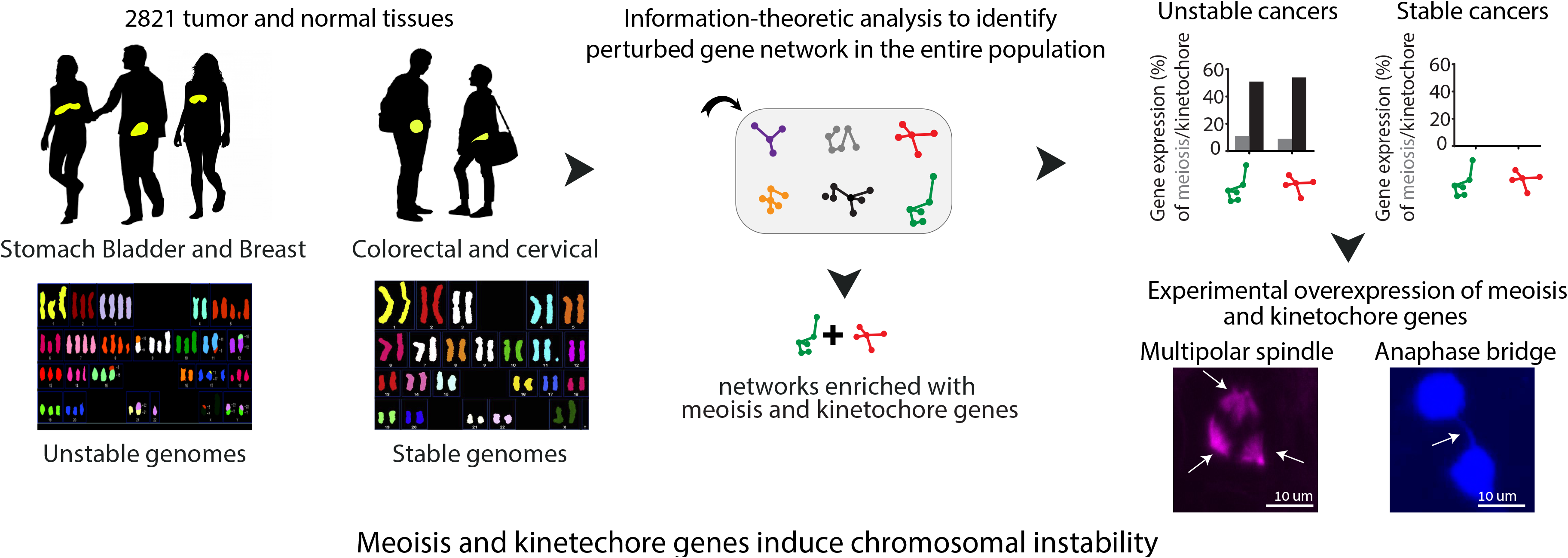
Schematic overview of how mis-expressed meiosis and kinetochore genes induce chromosome instability. 2821 tumors tissues of five cancer types obtained from TCGA datasets were analyzed. Cancer types were categorized into two subtypes: cancer with unstable genomes (unstable cancers) and cancer with a more stable genome (stable cancers). Information-theoretic analysis is utilized to study the altered gene expression networks in the entire population. We find that the most dominant cancer-specific altered networks in unstable cancers were enriched with meiotic and kinetochore but not in stable cancers. The experimental overexpression of meiosis and kinetochore genes in cancer cell lines induced genomic instability phenotypes: anaphase bridges (right) and spindle defects (multipolar spindle, left).

To further validate our hypothesis that meiosis and kinetochore genes drive CIN we performed experimental studies in genomically stable and unstable cancer cell lines (CIN+ and CIN,[8]). We have demonstrated that induced expression of representative meiosis and kinetochore genes in cancer cell-lines increases genome instability in this setting. Moreover, we show that this overexpression elevates significantly genome instability in genomically stable cancer cell lines, but less so in unstable cell lines. Overexpression of these genes also led to enhanced transformation and invasiveness properties of the cancer cell lines, providing experimental evidence for the involvement of meiosis and kinetochore genes in genome instability and cellular transformation. An overview of the study is summarized in Figure 1.

## Materials and Methods

### 1. Data analysis

#### 1.1 Thermodynamic-based information theoretical approach (Surprisal Analysis)

Matrix of gene expression data was obtained from TCGA database for each cancer type. Every dataset was profiled for thousands of transcripts (total 20,530). The matrix was used as an input for the information-theoretic surprisal analysis using MATLAB software [30] [31]. This type of analysis was utilized previously for the characterization of genomic/proteomic alterations and identification of molecular gene/protein correlation patterns characterizing big datasets [30,32,33]. Briefly, we identify the expected gene expression levels at the steady state (a state in which the biological processes are balanced), and deviations thereof for each transcript *i* in normal and tumor subsets. The deviations occur due to environmental/genomic constraints. Any biochemical/genetic perturbation can be considered as a constraint and elicit a coordinated change in a group of transcripts (subnetwork). These subnetworks are named **unbalanced processes** and are identified through calculations of G_*i*α_ values (=weights of participation) for each transcript *i* in each process α (α=1,2..3). **Table S1** lists G_*i*α_ values for all transcripts in each unbalanced in each cancer type. Each transcript can participate in more than one unbalanced process due to non-linearity of biological networks. Only the transcripts located on the tails of the distributions of G_*i*α_ values are analyzed further for biological meaning. Additionally, the analysis identifies an amplitude, λ_α_ (*k*), or an importance of each process α in each tissue *k* (**Fig. 3B,**). Plots of amplitudes for all unbalanced processes in breast and other cancer types can be found in **Table S2**.

Sign of G_*i*α_ and, λ_α_ (*k*) means correlation or anti-correlation between the transcripts in the same process α (in case of G_*i*α_) or α between the same processes in different tumors (in case of λ_α_ (*k*)). For example, if the process α is assigned the values: λ_α_ (1) = 37, λ_α_ (20) = 0, λ_α_ (33) = −39, it means that this process influences the tumors of the patients indexed 1 and 33 in the opposite directions, while it is inactive in patient 20. In order to calculate whether a particular transcript was induced or reduced due to a process α, the product G_*i*α_*λ_α_ (*k*) is calculated for each transcript. In summary, for each transcript we identify a set of unbalanced processes and quantify how important each process in each normal/cancer sample. Thus, a comprehensive map of unbalanced processes is obtained for each cancer or normal sample that allows to characterize each tissue in heterogeneous datasets in detail. Detailed description on how surprisal analysis is implemented in biology and how G_*i*α_ and, λ_α_ (*k*) are computed is provided in detail in [30,32,33]

#### 1.2 Computation of copy number variation

We obtained data of copy number variation (CNVs) in cancer population from the TCGA genomic database. Copy number variation (CNVs) are a type of structural variant involving alterations in the number of copies of specific regions of DNA, which can be either deleted or duplicated. These chromosomal deletions and duplications involve large stretches of DNA (that is, thousands of nucleotides, which may span many different genes) but can range considerably in size as well as prevalence. Only CNVs larger than 1 Mb (large CNVs usually correlate with the genomic stability) were considered for further analysis and are thus termed lCNVs (large chromosome number variations), but we also validated that similar results are obtained if other thresholds (0.5Mb and 2Mb) are implemented **(Fig. S1)**. The number of lCNVs for each sample were summed and determined using R tools. The distribution for each cancer type is shown in **Fig. 2**.

**Figure 2.**
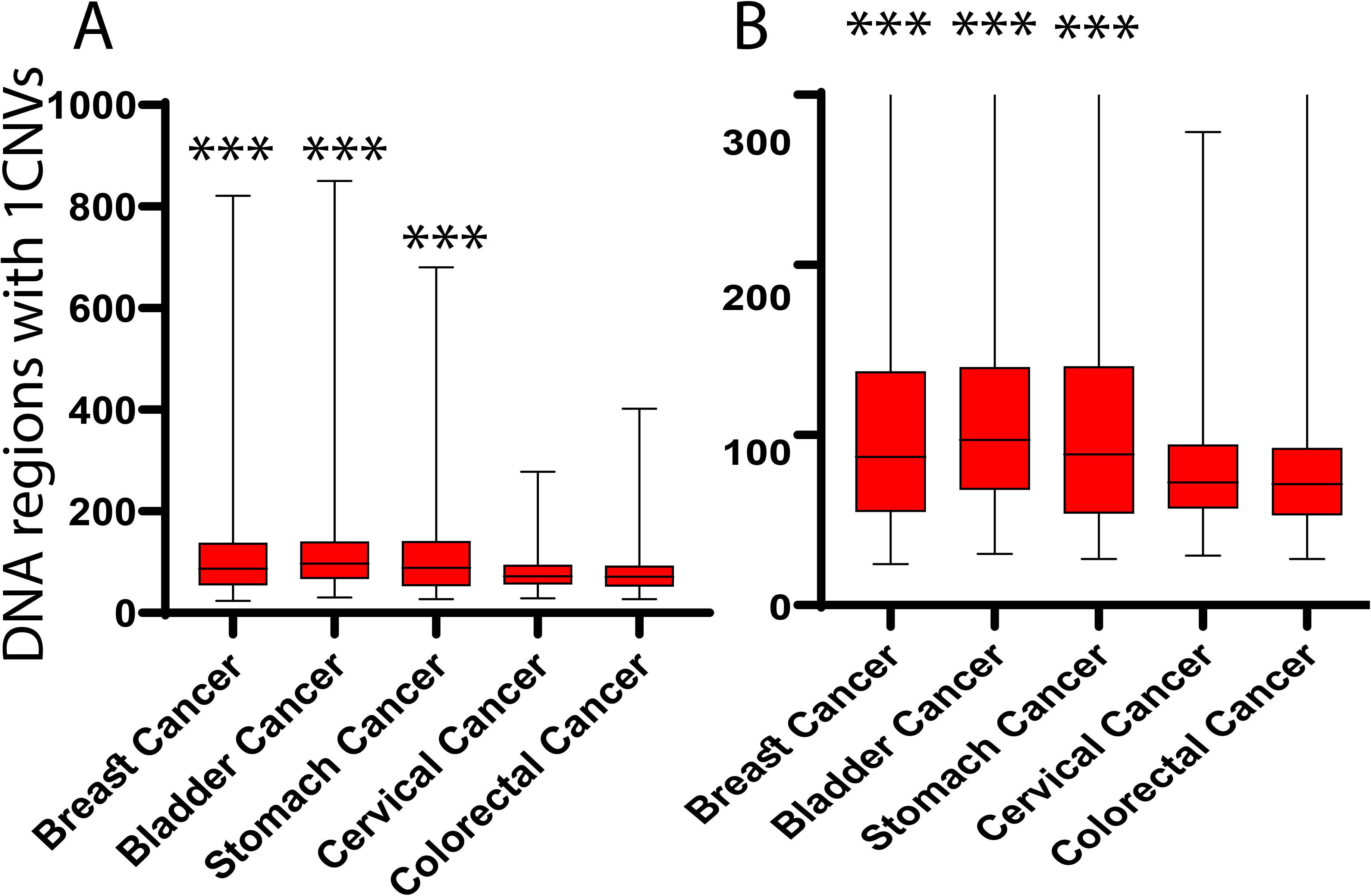
Distribution of Large Copy number variation (lCNVs) values in different cancer types. Bladder cancer has a high number of DNA regions with lCNVs (max 850, avg lCNVs= 112) followed by breast cancer and stomach cancers (max 821, avg lCNVs =109 and max 650 avg CNVs = 108 respectively). These three cancer types are categorized as unstable cancer types (p<0.001). On the other hand, colorectal, and cervical cancer types have a low number of DNA regions with lCNVs (max 402, avg lCNVs=79 and max 278, avg lCNVs= 80 respectively), and are categorized as more stable cancer types. Fig. 2A shows the full distribution of lCNVs values in all samples, while Fig. 2B shows a zoom–in on cases having up to 300 DNA regions with lCNVs (low CNV values).

### 2. Experimental methods

#### 2.1 Cell lines and culture

HCT116 human colon adenocarcinoma cell lines were maintained in Dulbecco’s Modified Eagle’s medium (DMEM, Sigma), supplemented with 10% fetal bovine serum (FBS), 1% PenStrep (100 U/mL Penicillin and 100 μg/mL Streptomycin) and 4 mM L-glutamine in a 37 °C incubator (5% CO2). MCF-7 breast cancer cell lines were maintained in RPM-1640, (Sigma), supplemented with 10% fetal bovine serum (FBS), 1% PenStrep (100 U/mL Penicillin and 100 μg/mL Streptomycin) and 4 mM L-glutamine in a 37 °C incubator (5% CO2).

#### 2.2 DNA preparation and Transfection

Meiotic and kinetochore genes tagged with Enhanced Green Fluorescent Protein (EGFP) (from BD Biosciences) were used. The GenElute, HP Plasmid Midiprep Kits (Bio Basic Inc, Canada). was used to isolate the plasmid. All cells were transfected 24h after initial plating. Transfections were performed using the Mirus transfection reagent (cat-81094967, Zotal, USA) with a 4:1 (transfection reagent: DNA) ratio. Transfection efficiency for HCT116 and MCF-7 cells was evaluated by counting the number of GFP positive cells by immunoflorescent microscope and calculating the percentage based on the total number of cells.

#### 2.3 Cell synchronization and immunostaining

HCT116 and MCF-7 cells were synchronized by double thymidine block [34]. Cells were treated with 2mM thymidine for 18 h in medium supplemented with 10% FBS. After washing twice with PBS, cells were cultured in fresh medium for 9h and again treated for 15h with media containing 2mM thymidine (10% FBS). After washing cells with PBS, the block was released by the incubation of cells in fresh medium and cells were harvested at 9h (HCT116) and 11h (MCF-7) and fixed with methanol. After that immunostaining was performed. Cells were washed 3 times with PBS and blocked with 5% BSA diluted with PBS. After that, cells were incubated with first Ab (α-tubulin antibodies, T5168, Sigma, USA, 1:400 dilution) for 2 hr at room temp. After washing twice with PBS, cells were incubated with a secondary Ab (donkey anti-mouse IgG antibodies, life science, USA, 1:200 dilution) for 1 hr at room temp followed by washing the cells twice with PBS. The cells were then incubated with Hoechst 33342 (cat: PIR-62249 Thermo scientific, Germany) diluted 1:10,000 for 5-30min for DNA visualization.

#### 2.4 Soft agar assay

Colony formation on soft agar was assayed in triplicate by plating 5000 cells in a layer of 0.3% (w/v) agar in assay DMEM (HCT116) and RPMI medium (MCF-7) medium, on top of a 0.6% (w/v) agar layer. Plates were incubated at 37 °C and 5% (v/v) CO2 for 3 weeks, and the medium was replaced every 4 d. Colonies were stained using 0.005% (w/v) Crystal Violet solution, and an image of the whole well was acquired using an Olympus SZ61 stereomicroscope. Colonies were counted using ImageJ software (http://imagej.nih.gov/ij/). The area and number of colonies was calculated.

### 3. Statistics

Significance was determined using a two-tailed Student’s *t* test. The p < .001 was considered as extremely significant (***), *p* < .01 as highly significant (**), and *p* < .05 as statistically significant (*).

## Results

### Degree of genome instability varies between different cancer types

We hypothesized that the extent of alterations in gene expression levels of meiotic and kinetochore genes may be related to the degree of genome instability. To explore this, we determined the distribution of large DNA tracts which exhibited copy number variations in five different cancer types, namely breast (n=1104), bladder (n=407), stomach (n=415), colorectal (n=383) and cervical (n=305) cancer patients and compared them to normal tissues from breast (n=114), bladder (n=19) stomach (n=20) colorectal (n=51) and cervix (n=3), all from TCGA database. Copy number variation (**CNV**) is a type of chromosomal structural variation that involves alterations in DNA copy number of specific genome regions. Those regions can be either deleted or duplicated. The chromosomal deletions and duplications can involve large stretches of DNA, e.g. thousands of nucleotides, which may span many different genes. In our analysis we included only CNVs that were larger than 1 Mb (lCNV= large CNVs of more than 1*10^6^ Base pairs, 1 Mb). The 1Mb cutoff was used as this size of CNV occurs rarely in normal human population (CNVs of >1 Mb occur naturally in <1% in the general population ([35]), but can be found in cancer tissues. Despite that, we have also analyzed a threshold of 0.5 Mb and 2 Mb with no change in the conclusions (**Fig S1**). We found that Bladder tumors had a large number of DNA regions with lCNVs (<u>high lCNV value</u>) with a maximum value of 850 lCNVs and an average value of 112 lCNVs. Breast and stomach cancers also show high values of lCNVs (with a maximum of 821 lCNVs, and average of 109 lCNVs for breast and a maximum of 650 lCNVs and average of 108 lCNVs for stomach cancer) (**Fig 2A,B**). On the other hand, colorectal and cervical cancer types have less DNA regions with lCNV (max 402, avg lCNVs=79 and max 278, avg lCNVs= 80 respectively) **(Fig. 2A, B)**. **(p<0.001)**. This finding allowed us to compare between low and high CNV cancers and confirm a relationship between altered expression of meiosis and kinetochore genes and genome instability.

### Altered gene expression networks, characterizing cancer samples with genome instability, are enriched with overexpressed kinetochore and meiosis transcripts

To explore the relationship between the degree of genome instability (high or low CNV) and the altered expression of meiosis and kinetochore we performed a large scale, unbiased computational analysis of over 2800 normal and cancer tissues from the 5 cancer types mentioned above. We utilized a computational information-theoretic surprisal analysis (<u>SA</u>) [32,33] in order to identify altered gene correlation co-expressed networks in cancer tissues, named *unbalanced processes*. Several different unbalanced processes (=subnetworks) may occur in a particular cancer type due to inter-patient heterogeneity **(Fig. 3A)**. SA deciphers a number of unbalanced processes in each cancer type, by calculating the expected expression levels of the tested molecules, such as transcripts or proteins, at the steady state (i.e. the balanced, unconstrained state), and the deviations thereof due to environmental or genomic constraints. These constraints elicit coordinated changes in expression levels of transcripts/proteins, named unbalanced processes (see [32,33] for more details). Co-varying altered transcripts that deviate from the steady state significantly and in a coordinated manner are grouped and each group represents an unbalanced process α. The analysis determines those transcripts through calculation of a “weight of participation”, G_*i*α_, of each transcript *i* in a process α (**Table S1**). Every altered transcript can be involved in several unbalanced processes.

Next the analysis assigns an amplitude, λ_α_ (*k*), an importance of each process α in each tissue *k* **(Fig. 3B)**. **Table S2** lists amplitudes of all processes in every cancer type and in every tissue. Several distinct unbalanced processes can be active in each cancer type/cancer tissue [33,36]. Detailed description of the analysis can be found in [33].

**Figure 3.**
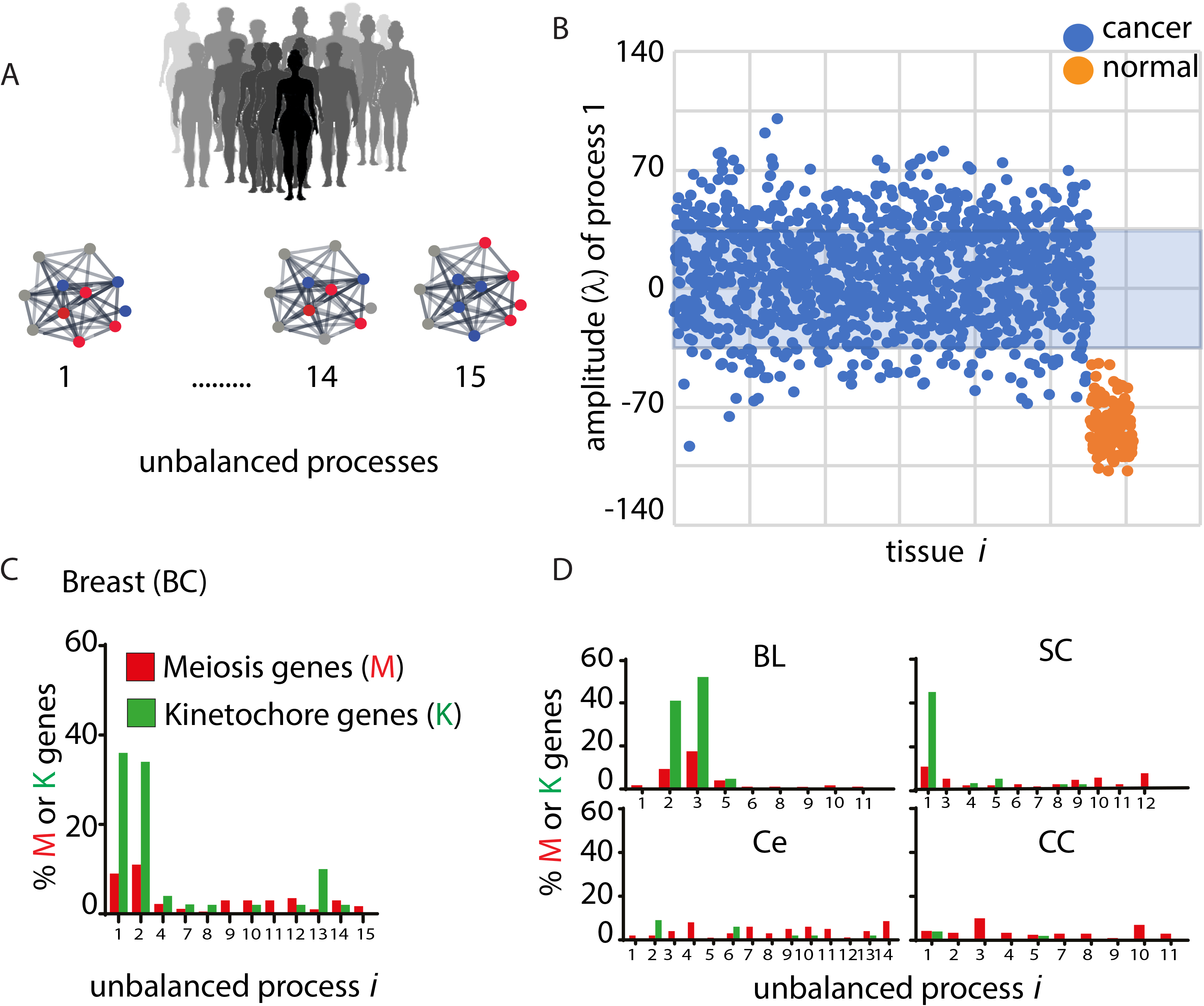
Involvement of kinetochore and meiosis genes in unbalanced processes as identified using surprisal analysis in both unstable and stable cancers. Every dataset is profiled for thousands of transcripts (total 20,530), which are resolved into altered networks (unbalanced processes) characterizing each tumor and normal tissue. All transcripts, which deviate from the balance state in the same (coordinated) way, are organized in groups, unbalanced processes (lower panel). **(B)** Amplitudes (λ_1_ (*k*)), representing an importance of a process α in each tissue, are shown for the most dominant process 1 in breast dataset. Tumor tissues are represented by blue dots and normal tissues by orange. This process clearly distinguishes between cancer (blue dots) and non-cancer (orange dots) tissues. For example, 21% of cancer tissues harbor this unbalanced processes (tissues with positive (λ_1_ (*k*) amplitudes). The blue box marks threshold limits. **(C)** Meiosis (red) and kinetochore (green) genes were found to participate in the most cancer-specific dominant processes in breast cancer: processes 1 and 2. **(D)** Meiosis (red) and kinetochore (green) genes were found to participate in the most cancer-specific dominant processes in bladder cancer: processes 2, 3 and 4 and stomach cancer: processes 1 and 3. There is a significantly lower percentage of meiosis and kinetochore genes in the dominant processes characterizing colorectal and cervical cancers (lower panel).

Using SA we identified 16 distinct unbalanced processes in breast cancer of which 12 were determined to be *cancer specific* **(Fig. 3C,** for example process 3 appears in both normal and cancer tissues, thus does not appear in the plot of **Fig. 3C)**. Rigorous error analysis, as described in [33,36,37] and Methods, was applied in order to determine a number of unbalanced processes beyond the noise in each dataset. Processes with lower indices, such as processes 1 and 2, were the most dominant and appeared in high a percentage of the patients. For example, the most dominant process, process 1, which was found in 21% of breast cancer patients (239 of 1095 breast cancer samples with positive (λ_1_ (*k*) amplitudes, **Fig. 3B;** see Table S3 which includes biological categories characterizing this and other processes. Genes with positive G_*i*α_ values (Tab “G1 positive” in Table S3) are induced in the tissues with positive λ_1_ (*k*) amplitudes and genes with negative G_*i*α_ values are reduced in those tissues and vice versa. See Methods for more details). Process 2 appeared in 20% of breast cancer patients **(Table S2).** Those processes were enriched for induced meiosis/ kinetochore genes (**Fig. 3C**). Interestingly, less common unbalanced processes (with higher indices) included significantly less kinetochore and meiosis genes **(Fig. 3C**). Similar results were found for the two other types of cancer with high values of lCNVs: bladder cancer and stomach cancer **(Fig. 3D,** upper panel**; Tables S4, S5** include all biological categories associated with those processes).

In contrast, the most dominant unbalanced processes (**Table S6-S7** include all biological categories associated with those processes) in cancers with lower values of lCNVs (colorectal and cervical cancers) were not particularly enriched for meiosis and/or kinetochore genes (**Fig. 3D,** lower panel). These results show a correlation between high lCNV values and overexpression of meiosis/ kinetochore genes **(Fig.2 and Fig.3 C,D)** and evince a possibility that highly expressed meiosis and kinetochore genes might be involved in genome instability of those cancers.

To further investigate the correlation between the altered expression levels of meiosis/kinetochore genes and genome instability, we examined every cancer type individually. We compared cancer samples with highly unstable genomes (10% of the samples with highest lCNVs values) to the cancer samples with relatively stable genomes (10% of the samples with lowest lCNVs values) within <u>the same cancer type</u>. The results demonstrate that in breast cancer, the most dominant processes 1 and 2, harboring a large number of induced meiosis/kinetochore genes, appeared in a relatively high percent of cancer tissues with high lCNVs (~25% and 50% respectively, **Fig. 4A**). Patients having more stable genomes (lower lCNVs) did not harbor those processes (**Fig. 4A**). Similar results were found in bladder and stomach cancers (**Fig. 4B-C).** In contrast, such a correlation between the enrichment of induced meiosis/kinetochore genes and genome instability was not found in the cancer types with relatively stable genomes (colorectal and cervical cancers, **Fig. S2)**.

**Figure 4.**
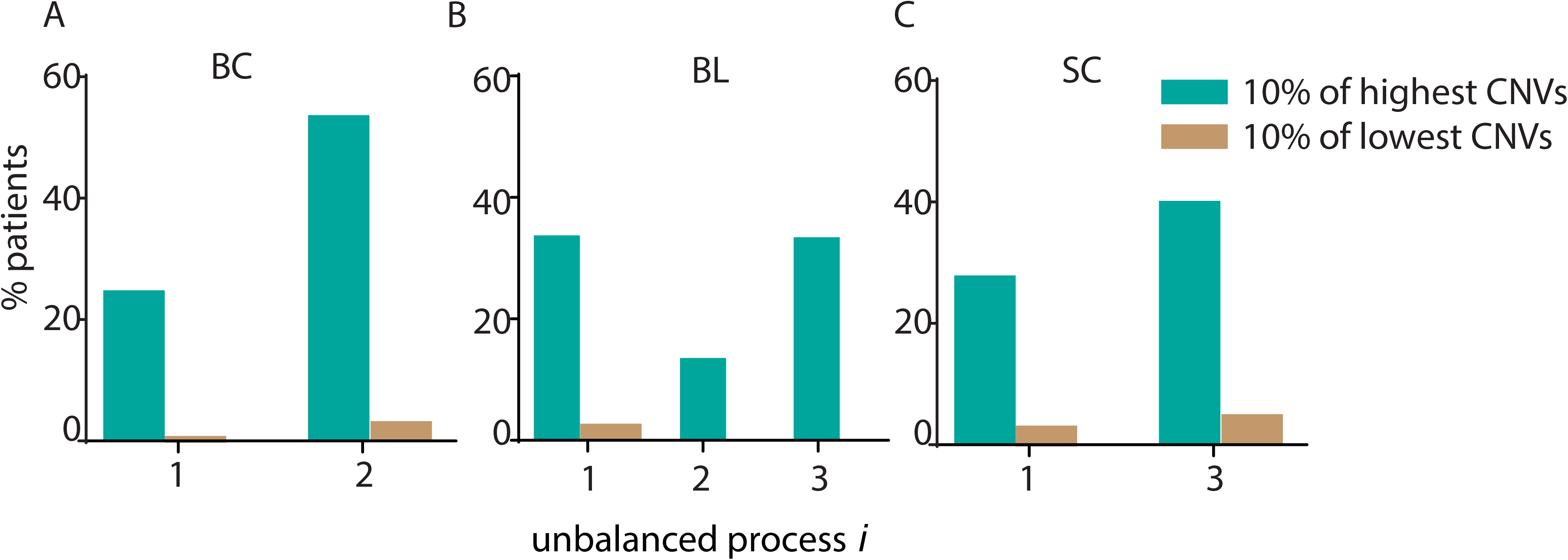
The correlation between over-expression of meiosis/kinetochore genes and genomic instability within each cancer type. Unstable breast (A), bladder (B) and stomach cancers (C) were analyzed separately as following: Distribution of lCNVs values was generated for each cancer type. 10% of the samples with highest lCNVs values (unstable group) and 10% of the samples with lowest lCNVs values (stable group) were selected in each cancer type. Cancer specific unbalanced processes of 10% of the samples with highest lCNVs values were compared to the processes appeared in 10% of the samples with lowest lCNVs values. The most dominant processes harboring meiosis/kinetochore genes appeared in more patients in the unstable group in all three cancer types.

These results point to a high correlation between the large and enriched groups of induced meiosis/kinetochore transcripts in dominant cancer-specific processes and genome instability. This suggests that meiosis and kinetochore genes may have an active role in driving CIN. This strong correlation prompted us to examine this hypothesis experimentally.

### Over expression of representative meiosis and kinetochore genes in genomically stable and unstable cancer cell lines

In order to test our hypothesis, we selected several representative kinetochore and meiosis genes participating in the dominant cancer-specific unbalanced processes (with high G_*i*α_) and overexpressed them in cancer cell lines. The most dominant kinetochore gene, which was found to be associated with CIN, was HJURP (found in the dominant unbalanced processes of all high CIN cancers, **Table S8**). HJURP (together with the protein it chaperones, CENP-A) represents the structural basis of the kinetochore structure [19,38].

CENP-A is the histone H3 homolog that forms a platform upon which all other kinetochore components assemble [39] [40]. Since CENP-A is functionally related to HJURP and was also found in all dominant processes of the high CIN tumors, we have also overexpressed CENP-A in our assays.

To select representatives among the meiotic genes we looked at the unbalanced process 3 in bladder cancer as it included the highest number of meiosis-related genes in comparison to other tumors analyzed. We thus overexpressed the two most dominant genes in this process: DMC1 and SMC1B.

As a negative control, we overexpressed *REC8*, which is a bone fide meiotic gene but was not found to participate in any dominant unbalanced processes in our analysis. REC8 is a meiosis specific component of cohesin, and participates in homologous chromosome pairing and in sister chromatid mono-orientation [41–43].

All genes were overexpressed in two cancer cell-lines-HCT116, a colon cancer cell line which is CIN negative and has a relatively stable genome, and MCF7, a breast cancer cell line which shows high chromosome instability [8,44]. The genes were fused to GFP to monitor their expression (**Fig. S3**).

### Over expression of meiosis and kinetochore genes promotes genome instability in cancer cell lines

In order to check whether the selected meiosis and kinetochore genes promote genome instability we evaluated the number of cells with lagging chromosomes, anaphase bridges, uneven segregation of chromosomes and deviation from a bipolar spindle configuration as a means to estimate genome stability [25,45–47]. **Figure 5 (B, C and G)** shows that overexpression of the kinetochore genes, CENP-A and HJURP, and one of the meiosis genes, DMC1 (but not SMC1B) in HCT116, caused a significant elevation in anaphase bridges and lagging chromosomes. Overexpression REC8, the negative control we used, did not affect the chromosome segregation phenotype. **Figure 5 (E, F and I)** also shows that overexpression of all meiosis and kinetochore genes in HCT116 cells caused a significant elevation in mono-polar and multi-polar spindle formation compared to an empty plasmid.

**Figure 5.**
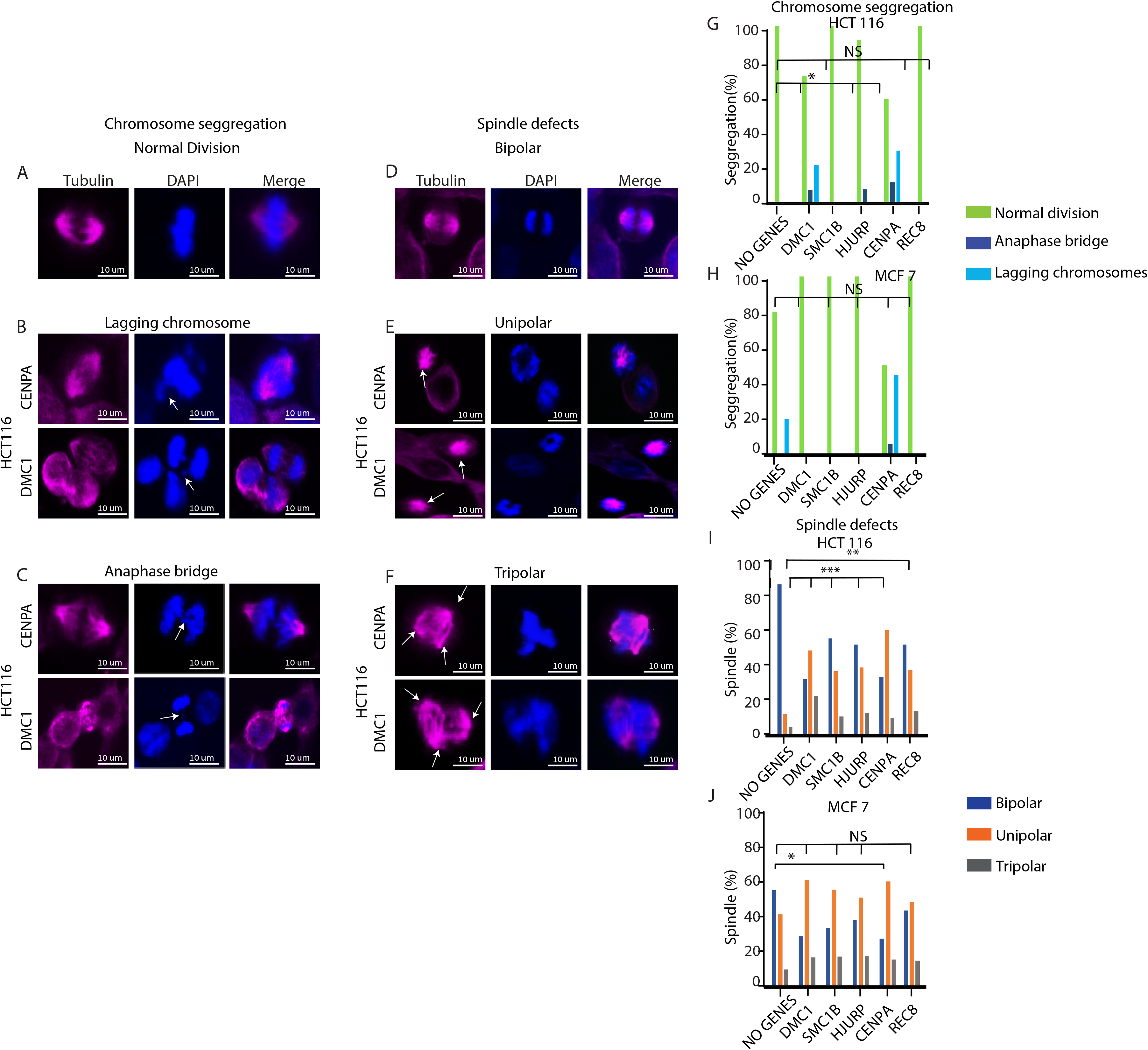
Overexpression of meiosis (DMC1 and SMC1B compared to REC8 as a control) and kinetochore genes (CENP-A and HJURP) causes a genomic instability phenotype. The effect of overexpressed meiosis and kinetochore genes in stable (HCT116) and unstable (MCF-7) cancer cell lines was measured by staining cells with DAPI and anti-tubulin IF (see Methods). Scoring the cells for genome instability was performed by counting defects during anaphase (anaphase bridges, lagging chromosomes) and apparent spindle formation defects (multipolar and unipolar). Presented are immunofluorescence images showing the effects of overexpression of meiosis (DMC1) and kinetochore (CENP-A) genes. Showing abnormal cell divisions i.e. Anaphase Bridge (**B**) and lagging chromosomes (**C**) and apparent spindle formation defects (unipolar **E** and tripolar **F**). Quantification of chromosome segregation and spindle defects in meiosis and kinetochore transfected stable cell line HCT116 shows more significant defects compared to the MCF 7 cell line (**G-J)**. Statistical significance is shown by asterisks (*P < 0.05; **P < 0.01; ***P < 0.001). Scale bar equals 10μm.

In contrast, overexpression of meiosis and kinetochore genes in the genomically unstable MCF7 cells caused a significantly less severe phenotype **(Figure 5, H and J**). Only CENP-A overexpression caused a small elevation in the occurrence of mono-polar and multi-polar spindles (**Fig. 5J**). However, all other phenotypes, related to the spindle (**Fig. 5J**) and chromosome segregation (**Fig. 5H**), were not significantly affected in MCF7 cells in response to induced expression of meiosis/kinetochore genes.

These results demonstrate that overexpression of our identified kinetochore and meiosis genes in genomically stable cells has the ability to promote genome instability. The same overexpression has a significantly smaller effect in a cell line that has already acquired a high degree of genome instability before the gene transfection.

### Over expression of meiosis and kinetochore genes promotes invasiveness of cancer cells

Cancers with unstable genomes are often more invasive than cancers with stable genomes [8,48,49]. Therefore, we hypothesized that overexpression of meiosis and kinetochore genes and promotion of genome instability could induce invasiveness and transformation properties of cancer cells.

To examine a change in the transformation properties of the cells we tested an ability of HCT116 and MCF7 cells to generate colonies in soft agar following overexpression of the meiosis/kinetochore genes [50,51]. **Figure 6 and S4** show that overexpression of all meiosis and kinetochore genes in HCT116 cells enhanced both the number of colonies generated and the size of the colonies, demonstrating enhanced cancer transformation properties. However, the number of colonies, overexpressing the negative control-Rec8 was smaller, although the area was similar to others. In general, the overexpression of kinetochore genes caused a greater effect than meiosis genes. In addition, overexpression of DMC1 created bigger colonies than the other meiotic genes we overexpressed. Surprisingly, although the genome instability parameters we previously checked were not increased in MCF7 cells upon the gene overexpression (see **Fig. 5**), an elevation in the number and size of the colonies in soft agar was detected in this cell line (**Fig. S5**). These results show that induced invasiveness and cellular transformation of the tested cell lines may correspond to the induced expression of meiosis and kinetochore genes, although not necessarily linked to the ability of those genes to induce genome instability parameters.

**Figure 6.**
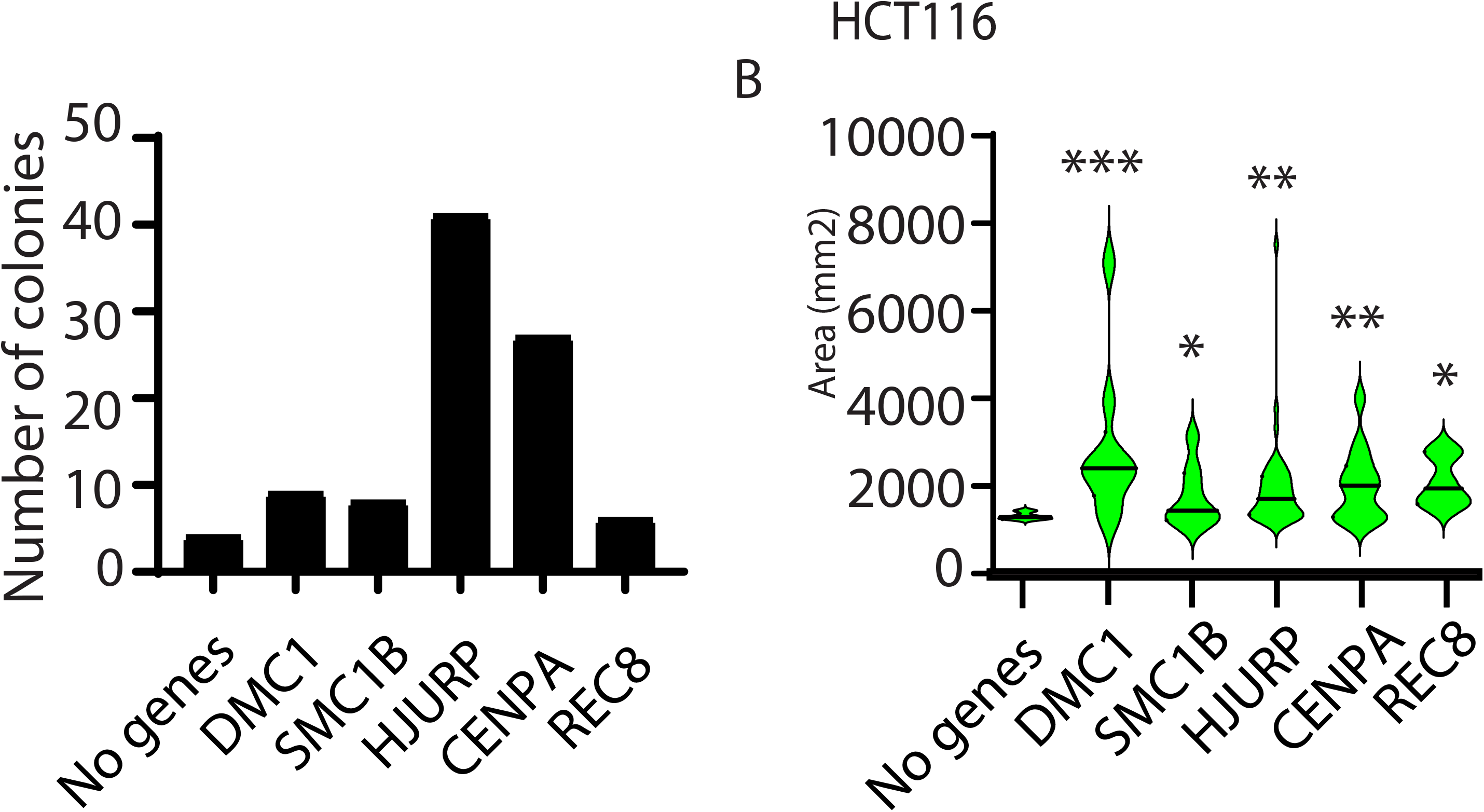
Overexpression of meiosis (DMC1and SMC1B) and kinetochore genes (CENP-A and HJURP) causes elevated invasiveness. Overexpressed meiotic and kinetochore genes in a stable cell line (HCT 116) have the ability to promote invasiveness in a soft agar assay. A soft agar colony formation assay was applied for the detection of transformed cells when overexpressed with meiotic and kinetochore genes. The number of colonies **(A)** and size of the colonies **(B)** were compared (see Methods). Statistically significant differences compared with empty vector (No genes) were determined using a Student’s 2-tailed t test (*P < 0.05; **P < 0.01; ***P < 0.001)

## Discussion

Genome instability and the mechanisms behind it are among central questions in cancer biology in recent decades [52]. Here we investigate a new possible route to achieve genome instability in cancer by the overexpression of genes which participate in meiosis or in kinetochore formation. The process of meiosis includes inherent genome instability which occurs through meiotic homologous recombination and sister chromatid mono-orientation. The perturbed expression of kinetochore proteins has also potential to affect the processes involved in proper chromosome segregation and genome stability.

We have taken a multi-pronged approach to support our hypothesis and have used computational analysis of large cancer datasets and an experimental approach utilizing cancer cell line models. Using information-theoretic surprisal analysis of five different cancer types, obtained from TCGA database, we have shown that the most abundant altered gene expression networks, characterizing unstable cancers, were enriched with meiosis and kinetochore transcripts. Altered gene expression networks, characterizing cancers with low lCNVs were not enriched with those transcripts. Moreover, in unstable cancers, patients with the highest lCNVs were characterized by the unbalanced processes enriched with meiosis and kinetochore genes, in contrast to the patients with low lCNVs within the same cancer type, which did harbor those processes.

Although these analyses were merely restricted to a correlation, the finding of this correlation in five major cancer types, and the extension of the correlation to the specific patient groups within each cancer hints to a strong link between the overexpression of meiosis and kinetochore genes and genomic instability in tumors.

In order to go beyond this correlation and demonstrate a causative effect, we performed experiments in which several representative meiosis and kinetochore genes, as identified by our computational analysis, were expressed in two cancer cell lines. The overexpression caused an elevation of genome instability parameters in the stable HCT116 cell lines, but less so in the unstable MCF7 cell line. However, overexpression caused both cell lines to increase their invasiveness and transformation properties as measured by colony formation ability in soft agar. These results could suggest that induced invasiveness (observed in both cell lines) is not directly related to the induced genome instability (observed in HCT116) although they both result from the overexpression of the same genes. Another possibility is that even a slight and undetectable increase in genome instability (as in MCF-7), can cause a large effect on the invasiveness of those cells. Further experiments are needed to distinguish between these possibilities.

Our results also demonstrate that meiosis and kinetochore genes can serve as markers for genome instability. Future work should assess the accuracy and sensitivity of those markers and whether downregulating meiosis and kinetochore genes could be used as a therapeutic approach.

In conclusion, we have shown that genome instability in tumors could be driven by overexpression of specific classes of genes, namely meiosis and kinetochore genes, which are involved in genome organization and maintenance of undifferentiated cells. This finding may have medical implications regarding the identification of genome instability in tumors, diagnosis and eventually the treatment of unstable tumors through manipulation of these gene networks.

---

**Figure S1.**
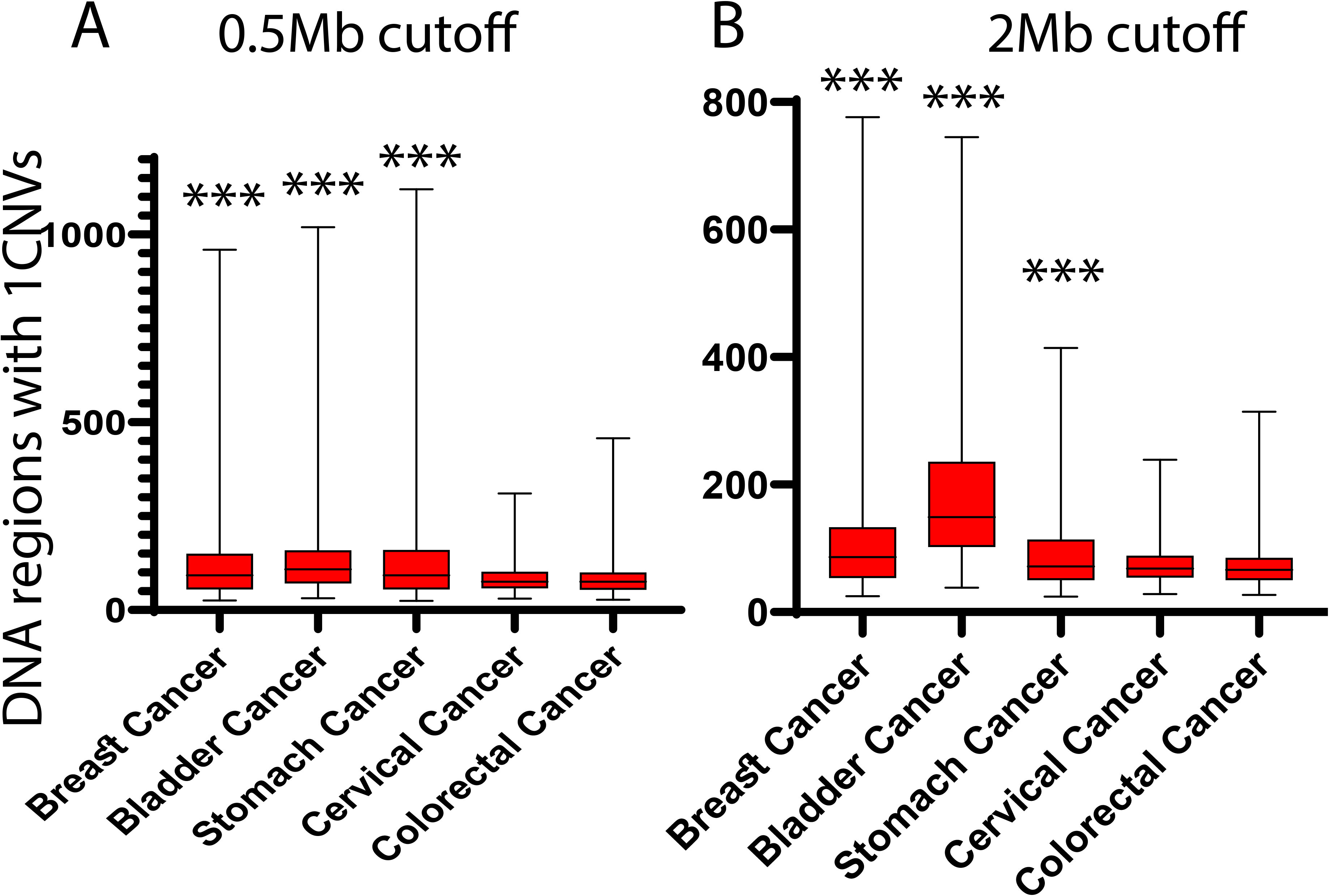
Distribution of Large Copy number variation (lCNVs) values in different cancer types in additional cutoffs (0.5Mb and 2Mb). The distribution of lCNVs in the different cancer types in two additional cutoffs: lCNVs larger than 0.5Mb **(A)** and larger than 2Mb **(B)**. The analysis of these threshold did not change the classification of the tumors into two disctict groups with high and low amounts of lCNVs.

**Figure S2.**
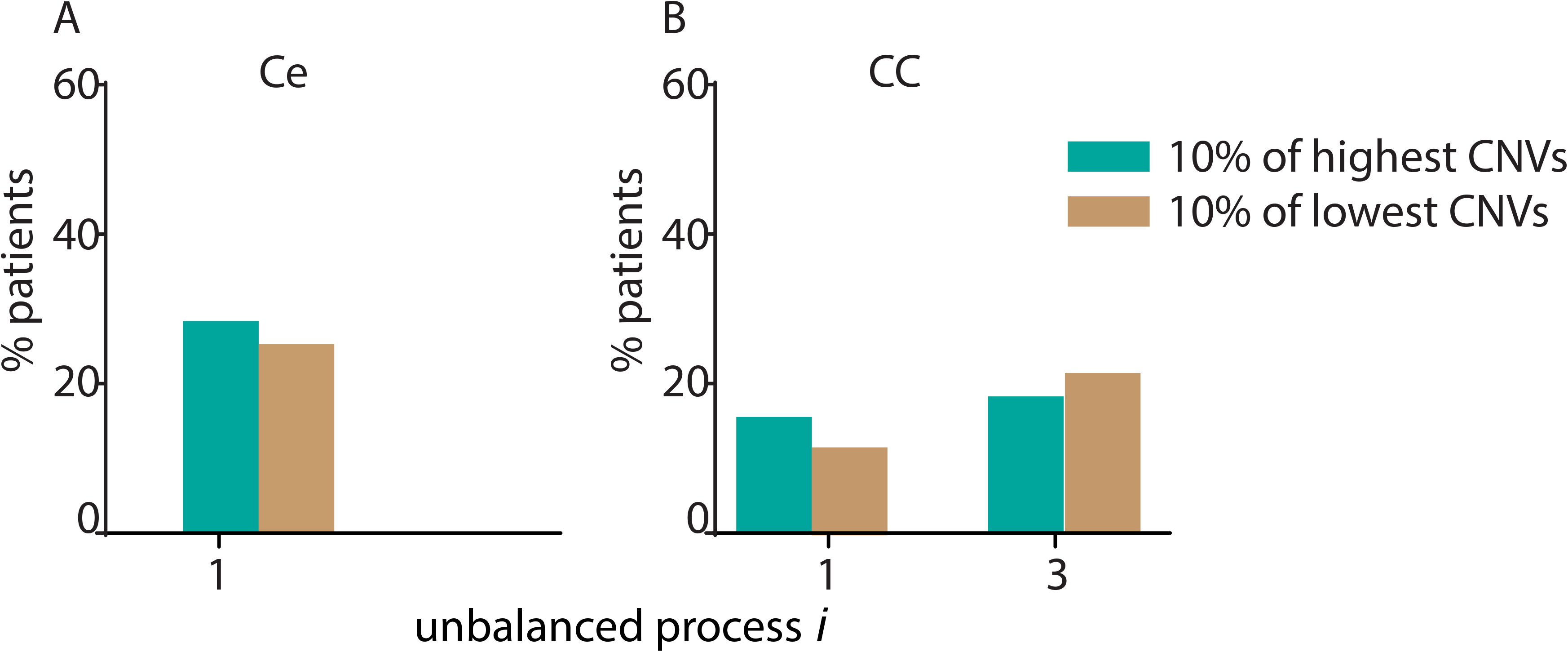
The correlation between over-expression of meiosis/kinetochore genes and genomic instability within each cancer type (stable cancers: cervical cancer (A) and colorectal cancer (B)). Cancers that more genomically stable (lower lCNV values) cervical and colorectal cancers were analyzed separately as following: Distribution of lCNVs values was generated for each cancer type. 10% of the samples with highest lCNVs values (unstable group) and 10% of the samples with lowest lCNVs values (stable group) were selected in each cancer type. Cancer specific unbalanced processes of 10% of the samples with highest lCNVs values were compared to the processes appeared in 10% of the samples with lowest lCNVs values. The most dominant processes appeared in the same percentage of patients in the stable and unstable group in the two cancer types.

**Figure S3:**
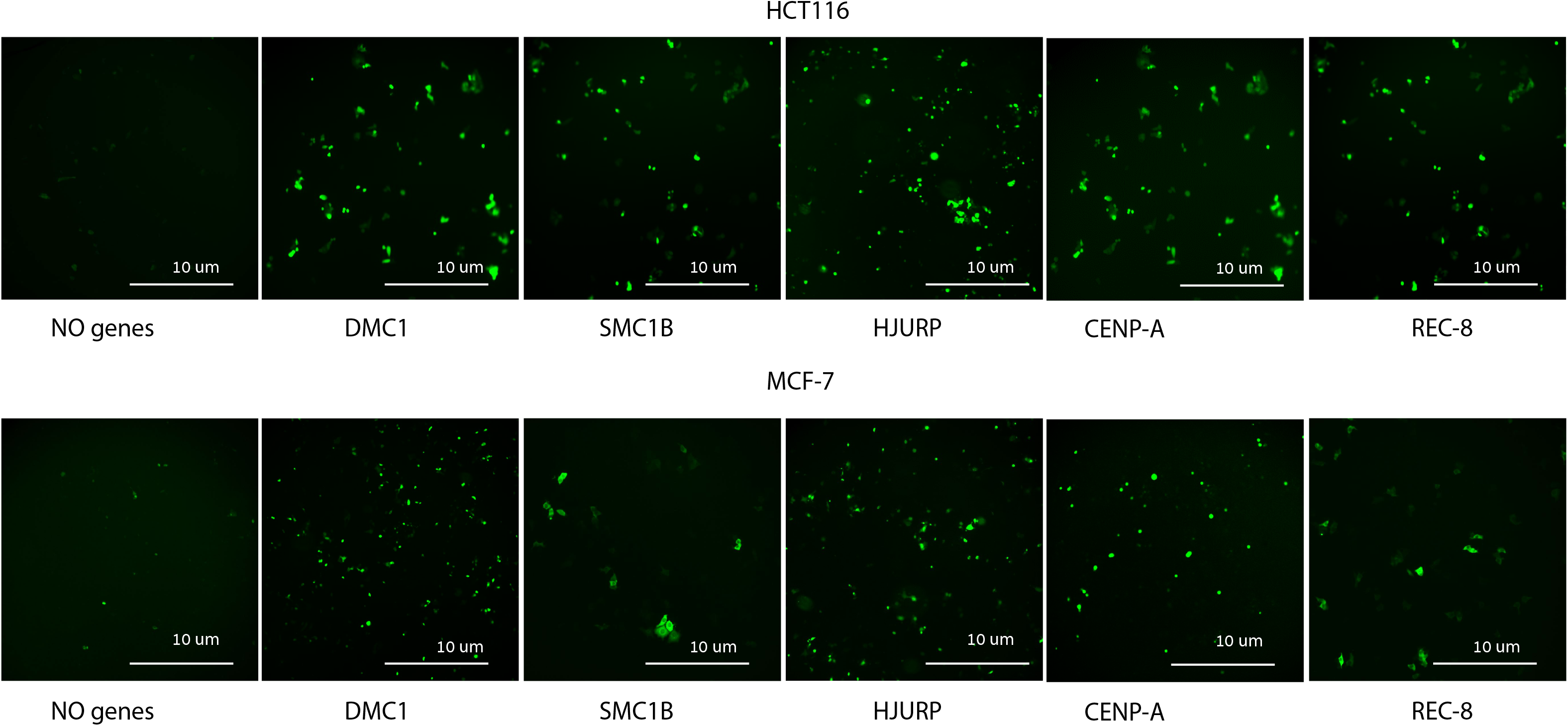
Over expression of meiosis and kinetochore genes: Meiosis/kinetochore genes were fused to GFP and the over-expression pattern in the cell lines shows that the overexpression was successful. Overexpression of meiotic genes (DMC1, SMC1B, REC8)and kinetochore genes (CENP-A and HJURP) in the HCT 116 cell line and MCF-7 cell line is presented.

**Figure S4:**
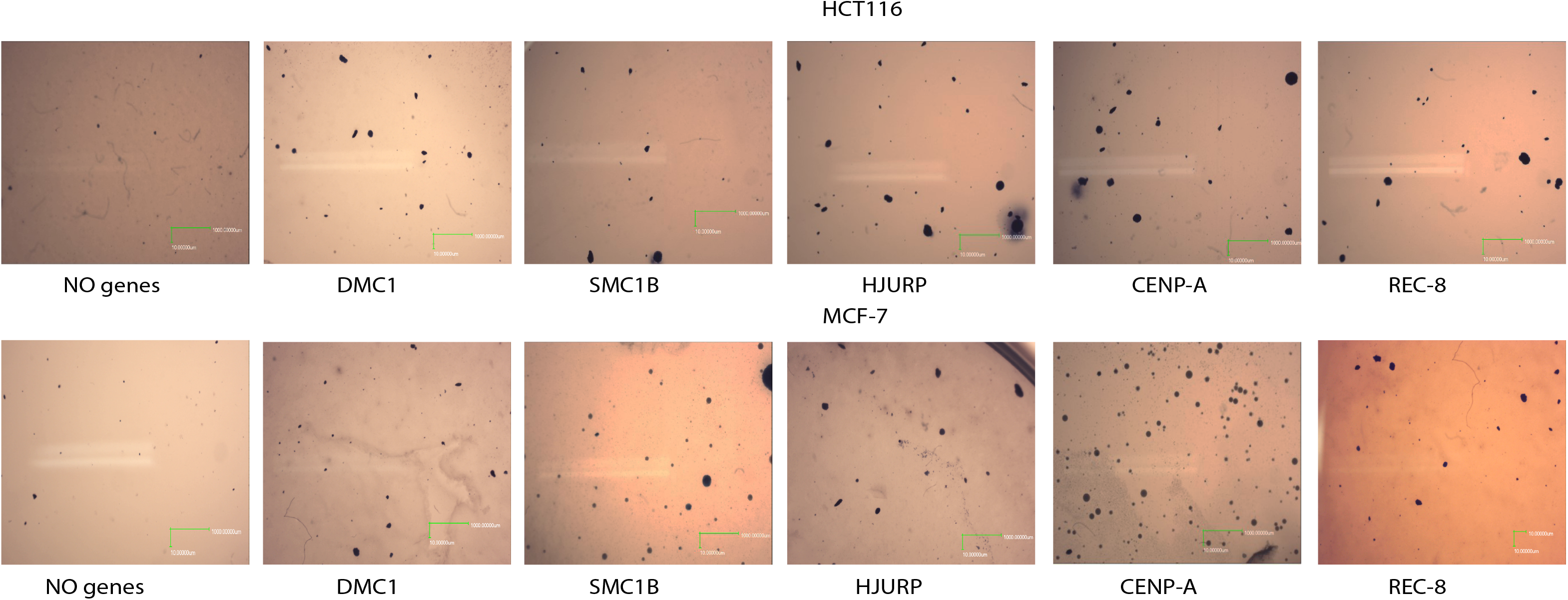
Meiosis/kinetochore genes induce number and size of colonies in soft agar. HCT116 and MCF7 cells overexpressing the meiosis/kinetochore genes mentioned above were seeded in soft agar. Representative images of soft agar assay for HCT116 and MCF-7 cell lines are shown.

**Figure S5:**
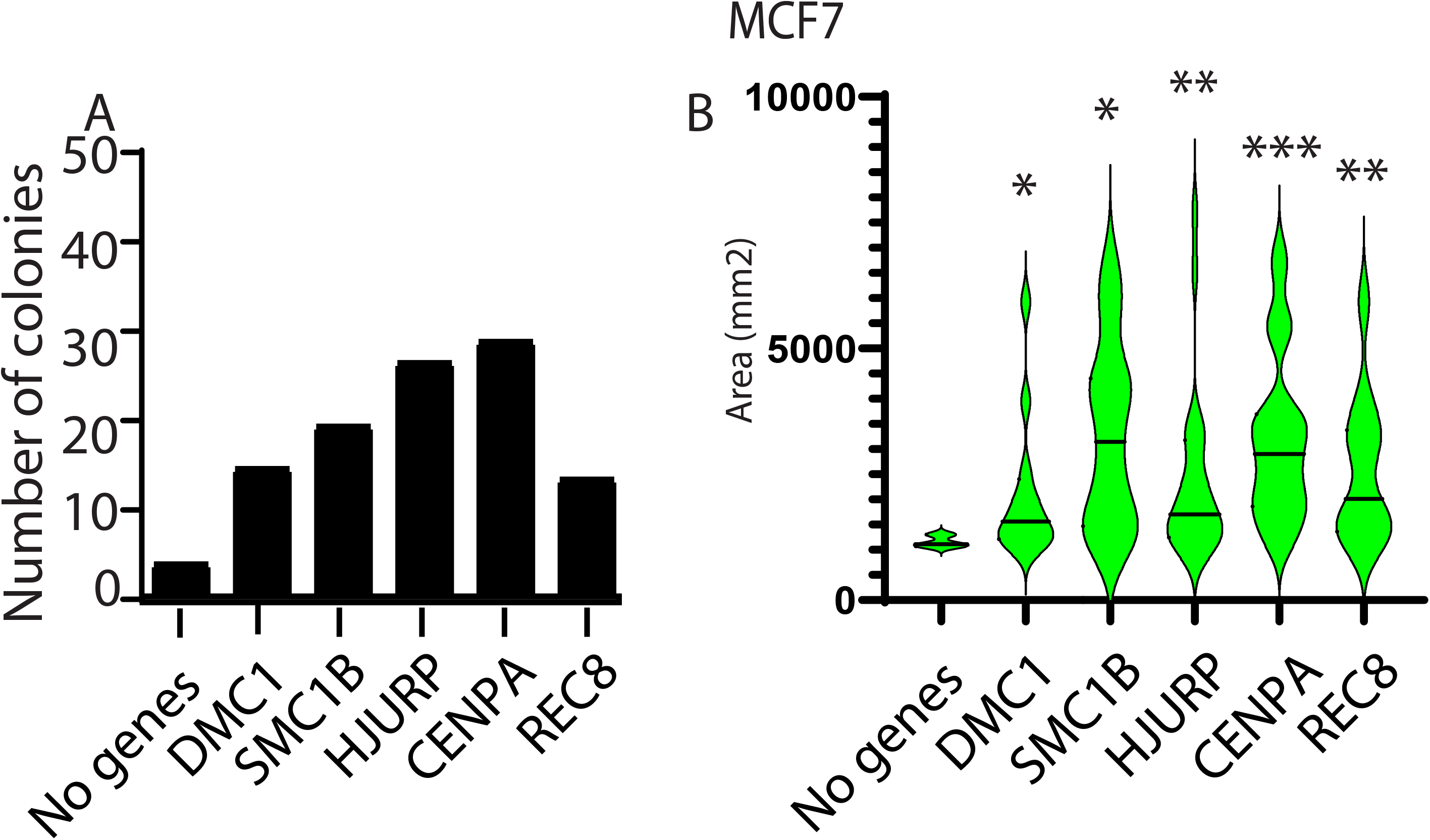
A soft agar colony formation assay was applied for the detection of transformed cells when overexpressed with meiotic and kinetochore genes in an unstable cell line (MCF-7). The number of colonies **(Fig A)** and size of the colonies **(Fig B)** were measured using imageJ. Statistically significant differences compared with empty vector (No genes) were determined using a Student’s 2-tailed t test (*P < 0.05; **P < 0.01; ***P < 0.001).

## Acknowledgments

We thank the Klutstein and Kravchenko-Balasha labs for fruitful discussions. This work was supported by a research grant from the Israeli Cancer Association (grant number 20190056) to MK and NKB; a special donation from the ICA USA section and Israel Science Foundation to NKB.

## Availability of data and reagents

The authors affirm that all data necessary for confirming the conclusions of this article are represented fully within the article and its tables and figures. All the necessary reagents from this study can be shared.

